# Mitochondrial defects leading to arrested spermatogenesis and ferroptosis in a mouse model of Leigh Syndrome

**DOI:** 10.1101/2022.11.22.517461

**Authors:** Enrico Radaelli, Charles-Antoine Assenmacher, Esha Banerjee, Florence Manero, Salim Khiati, Anais Girona, Guillermo Lopez-Lluch, Placido Navas, Marco Spinazzi

## Abstract

Impaired spermatogenesis and male infertility are common manifestations of mitochondrial diseases, but the underlying mechanisms are unclear. Here we show that mice deficient for PARL, the mitochondrial rhomboid protease, a recently reported model of Leigh syndrome, develop postpubertal testicular atrophy caused by arrested spermatogenesis and germ cell death independently of neurodegeneration. Genetic modifications of PINK1, PGAM5, and TTC19, three major substrates of PARL with important roles in mitochondrial homeostasis, do not reproduce or modify this phenotype. PARL deficiency in testis mitochondria leads to severe mitochondrial electron transfer chain defects, alterations in Coenzyme Q biosynthesis and redox status, and abrogates GPX4 expression specifically in spermatocytes leading to massive ferroptosis, an iron-dependent regulated cell death modality characterized by uncontrolled lipid peroxidation. Thus, mitochondrial defects can initiate ferroptosis *in vivo* in specific cell types by simultaneous effects on GPX4 and Coenzyme Q. These results highlight the importance of ferroptosis and cell-type specific downstream responses to mitochondrial deficits with respect to specific manifestations of mitochondrial diseases.

## INTRODUCTION

Impaired spermatogenesis and consequent infertility are increasingly common medical issues affecting about 9% of the global male population^1^. Oxidative stress and mitochondrial dysfunction are considered crucial pathophysiological causes, but their precise role is poorly characterized^2^. Moreover, male infertility has been reported to be a prevalent manifestation of mitochondrial diseases, although still largely unexplored^3^. Mitochondria play essential, but incompletely understood, roles in reproductive biology including spermatogenesis^4,5^. Mitochondrial diseases encompass various devastating inborn errors of metabolism caused by genetic defects in either mitochondrial or nuclear genome. How these gene defects most typically affect only specific organs/tissue types is currently not understood. This selective tissue vulnerability most likely involves cell-type specific activation of downstream molecular pathways that act independently or in parallel with the mitochondrial respiratory chain defects. Energy insufficiency alone does not provide a comprehensive explanation for the resulting clinical manifestations^6^. In this context, complex molecular responses to mitochondrial dysfunction are increasingly recognized as crucial homeostatic mechanisms ^7–9^

In our previous study we described PARL-deficient mice as a novel mouse model of Leigh syndrome^10^, one of the most common and severe mitochondrial diseases. PARL is an evolutionary conserved protease belonging to the rhomboid family inserted in the inner mitochondrial membrane with fundamental, yet controversial roles in cell homeostasis and human disorders such as Parkinson’s disease, Leber hereditary optic neuropathy, and type 2 diabetes^11–14^. A crucial role of PARL in mitochondrial fitness has been established thanks to seminal studies identifying its substrates^15–17^. These include, among others, PINK1, a mitochondrial kinase implicated in autosomal recessive forms of Parkinson’s disease and mitophagy^18,19^, PGAM5, a mitochondrial phosphatase implicated in Parkinsonism in mice^20^, and TTC19, a mitochondrial protein involved in the maintenance of Complex III activity and in human cases of Leigh syndrome^21,22^.

Here we show that impaired spermatogenesis represents the earliest phenotype of PARL*-*deficient male mice, preceding and independent of neurodegeneration. PARL deficiency leads to severe functional and structural mitochondrial abnormalities in germ cell mitochondria resulting in arrested spermatogenesis and induction of ferroptosis specifically in spermatocytes.

## RESULTS

### PARL deficiency results in arrested spermatogenesis and severe testis atrophy

PARL-deficient mice appear clinically normal until the age of 5 weeks, but they succumb by the age of 8 weeks because of the rapid development of a subacute necrotizing encephalomyelopathy that resembles Leigh syndrome^10^. Macroscopic reduction in testicular size is invariably present in postpuberal *Parl^-/-^* mice (Fig. 1A), confirming previous observations^10,23^, although we did not observe cryptorchidism^23^. At 5 weeks of age, before the onset of neurological manifestations, the testis weight of *Parl^-/-^* mice is almost half of that of matched WT littermates (Fig 1A). This striking difference is not explained by concomitant body weight reduction (Fig 1A). When compared to WT littermates, seminiferous tubules and epididymis from *Parl^-/-^* testis appear smaller in diameter with a complete lack of spermatids and spermatozoa (Fig. 1 B). Moreover, affected seminiferous tubules in *Parl^-/-^* testis are entirely populated by immature germ cells showing degenerative changes and prominent intraluminal exfoliation often in the form of multinucleated syncytia (Fig. 1B). Immunohistochemistry (IHC) for synaptonemal complex protein 1, SYCP-1, specifically expressed in primary spermatocytes during the zygotene and pachytene stage of prophase I of meiosis I^24^, and allograft inflammatory factor-1, AIF-1, specifically expressed in spermatids^25^, confirms that PARL deficiency causes complete meiotic arrest with seminiferous tubules entirely populated by prophase I primary spermatocytes (Fig. 1B). Distribution and morphology of other cell types in the seminiferous tubules and surrounding interstitium, including Leydig and Sertoli cells, appear normal.

**Figure 1.**
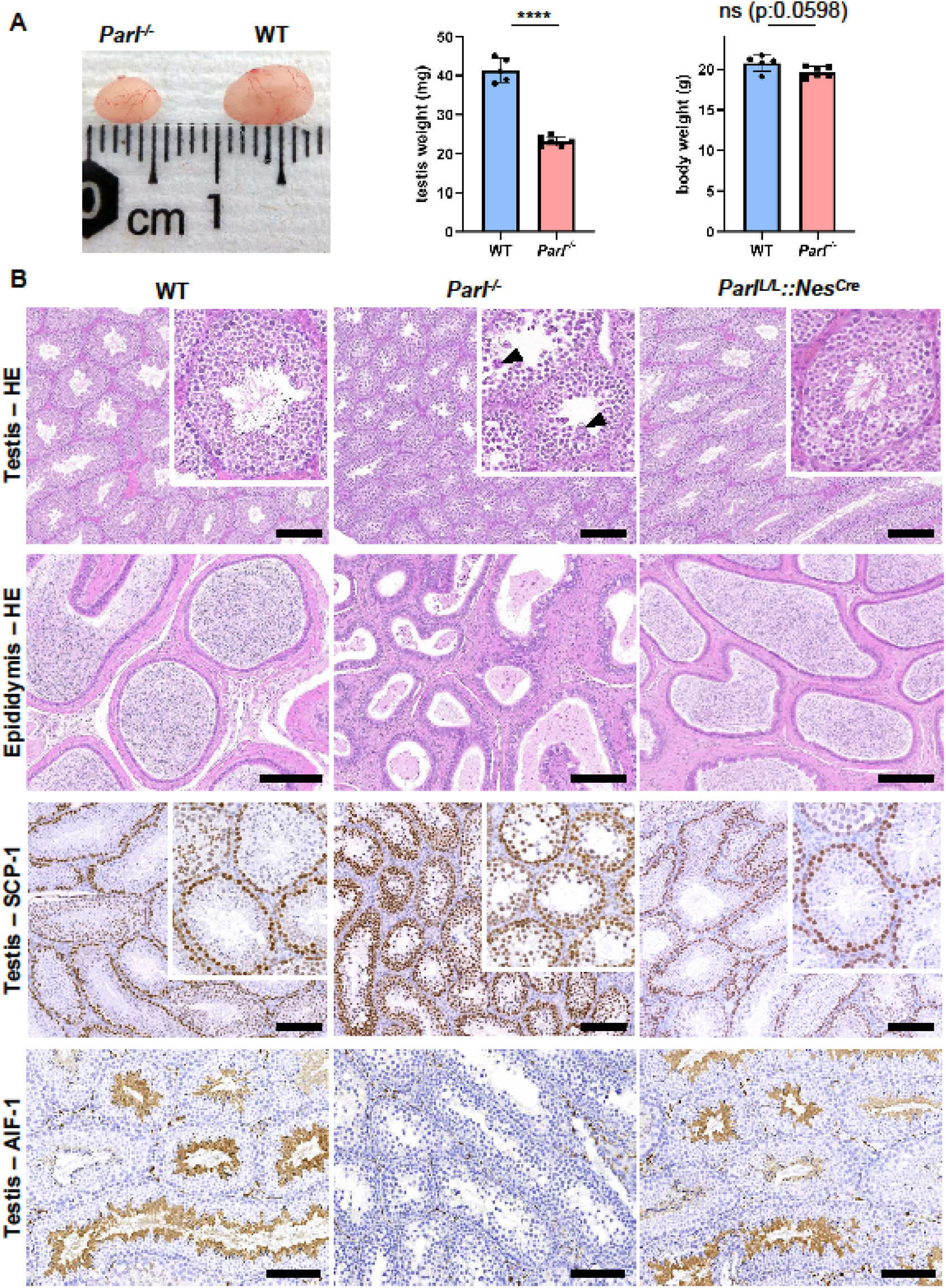
Severe testis atrophy in *Parl^-/-^* mice is caused by arrested spermatogenesis, independent from neurodegeneration. (A) Reduced testicular size and weight in 5-week-old *Parl^-/-^* mice (n=6) compared to WT littermates (n=5). The reduction in testicular weight is not explained by body weight differences. Unpaired two-tailed *t*-test, P-value < 0.0001. (B) Histological assessment of testes from postpubertal *Parl^-/-^* and WT mice at 6 weeks of age reveals reduced diameter of *Parl^-/-^* seminiferous tubules with impaired germ cell maturation and complete spermatogenesis arrest at the level of primary (premeiotic) spermatocytes (testis HE stain, n=10). *Parl^-/-^* seminiferous tubules also exhibits intraluminal exfoliation of degenerated spermatocytes often in the form of multinucleated syncytia (testis HE stain inset, arrowheads). The complete absence of sperm in *Parl^-/-^* epididymis compared to WT littermates demonstrates a complete arrest of spermatogenesis with no production of mature gametes (epididymis HE stain, n=10). Immunohistochemistry for synaptonemal complex protein 1, SCP-1, confirms complete spermatogenesis arrest at the level primary spermatocytes in *Parl^-/-^* testis (testis SCP-1, n=10). Unlike in *Parl^-/-^* testis, the distribution of SCP-1 expression in WT seminiferous tubules is confined to primary spermatocytes and it is lost in postmeiotic germ cells as they undergo maturation. Immunohistochemistry for allograft inflammatory protein 1, AIF-1, reveals the complete absence of spermatids in *Parl^-/-^* testis while WT seminiferous tubules are densely populated by AIF-1-positive spermatids at different levels of maturation (testis AIF-1, n=10). Mice with conditional *Parl* deletion driven by the *Nes* promoter in the nervous system and Leydig cells (*Parl ^L/L^::Nes^Cre^*) display a normal testicular and epididymal histology as well as SCP-1 and AIF-1 immunohistochemistry comparable to WT mice (right column, 8 weeks, n=4). Scale bars, 200 μm.

To rule out the contribution of the remote effect of subclinical neurodegeneration to the development of impaired spermatogenesis, we investigated whether mice with conditional deletion of *Parl* in the nervous system reproduced the same testicular abnormalities observed in the germline knockout. Eight-week-old *Parl^L/L^::Nes^Cre^* mice are affected by severe Leigh-like encephalopathy but show normal testicular size and histology comparable to WT littermates (Fig. 1B), indicating that the testicular disorder is not the consequence of neurodegeneration. As previously reported^26–28^, *Nes* is also expressed in Leydig cells (Figure 1-figure supplement 1) suggesting that the spermatogenetic defect is not secondary to PARL deficiency in these cells. Altogether, deficiency of PARL leads to male infertility via a complete arrest of spermatogenesis at the level of primary spermatocytes, independently of the effects of PARL in the nervous system and Leydig cells.

### PARL deficiency results in mitochondrial ultrastructural abnormalities and progressive degeneration and death of arrested spermatocytes

Next, we wondered whether the spermatogenesis defect induced by PARL deficiency is associated with pathological effects on mitochondrial morphology or with other features of cell degeneration and death. To answer this question, we performed a detailed morphological analysis using semithin sections and electron microscopy. In unaffected WT animals, germ cells progressively differentiate to spermatozoa following a maturation wave characterized by less differentiated germ cell forms (i.e., spermatogonia and spermatocytes) in the abluminal layers, more differentiated spermatids in the adluminal compartment, and mature spermatozoa in the lumen (Fig. 2A). Conversely, the analysis of postpuberal *Parl^-/-^* mice confirms impaired spermatogenesis showing severe vacuolar degeneration of arrested spermatocytes culminating in cell death with a clear severity progression from the abluminal to the adluminal compartment (Fig. 2A and 2B). Next, we assessed whether mitochondrial morphology was affected in PARL-deficient spermatocytes. Since differentiation *per se* leads to important morphological adaptations of mitochondria which parallel increasing bioenergetic demands requiring a shift from more glycolytic to more oxidative metabolism^29^, we focused our ultrastructural analysis on germ cells at the same stage of spermatogenic maturation.

**Figure 2.**
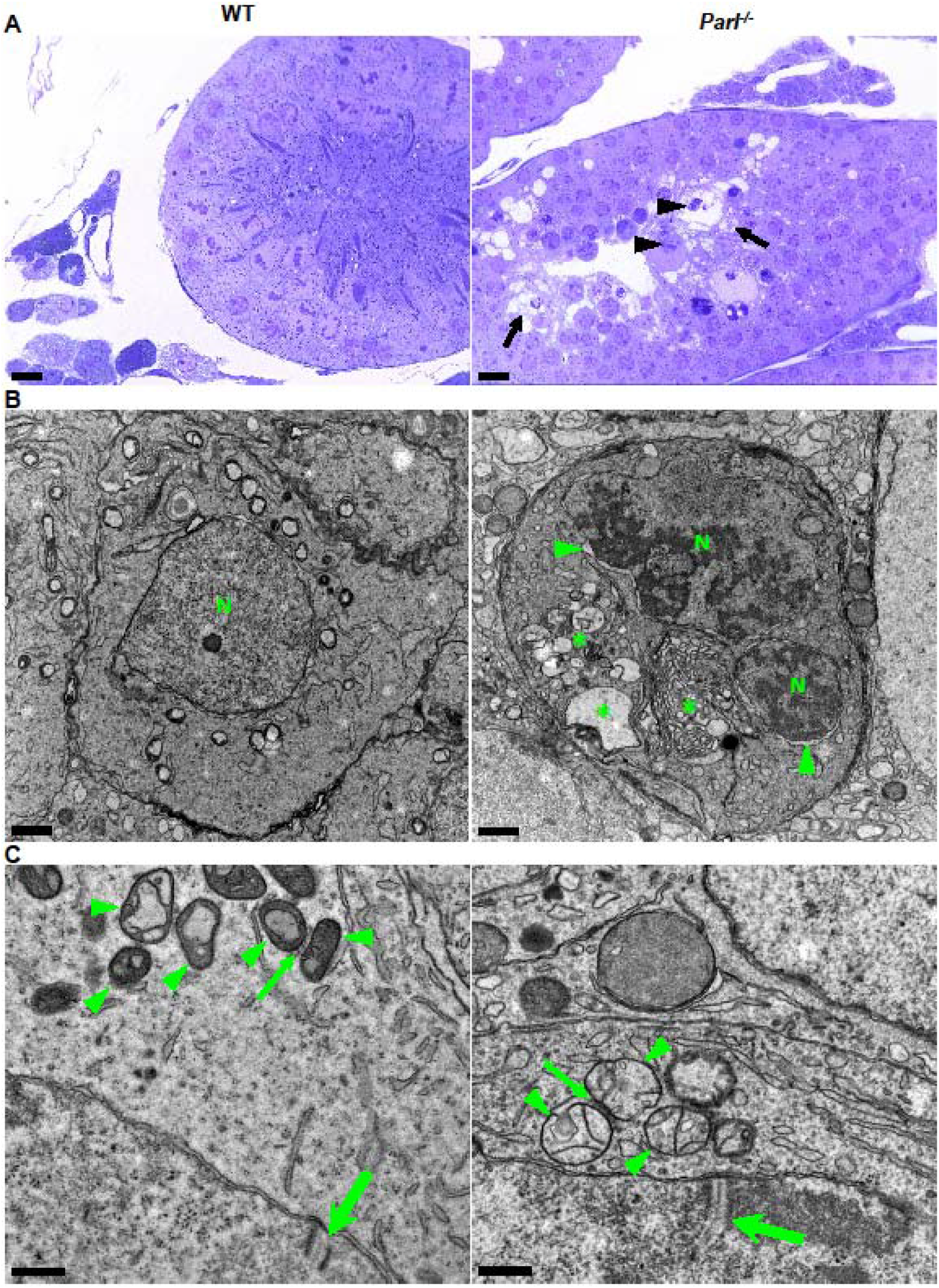
Impaired spermatogenesis in *Parl^-/-^* testis is associated with early mitochondrial morphological abnormalities and progressive degeneration of arrested spermatocytes. (A) Assessment of toluidine blue-stained semithin sections of testis from 5-week-old postpubertal WT and *Parl^-/-^* mice. Seminiferous tubules from *Parl^-/-^ mice* show extensive degenerative changes in arrested spermatocytes including tortuous membrane infoldings, cytoplasmic vacuolation (arrows), irregular chromatin clumping, nuclear fragmentation (arrowheads), and absence of germ cell maturation including spermatids and adluminal spermatozoa (5 weeks, n=3). A WT seminiferous tubule with normal germ cell maturation is shown for comparison (left panel). Scale bars, 20 μm. (B) Electron microscopy examination shows multifocal cisternae distention, disruption of the endoplasmic reticulum and Golgi apparatus, abundant accumulation of damaged membranous material and organelles (asterisks), and nuclear damage in *Parl^-/-^* spermatocytes. The nuclear envelope is diffusely distended (arrowheads) outlining a convoluted fragmented nucleus (N) with dense irregular clumps of chromatin. A WT spermatocyte at the end of pachytene is shown for comparison (left panel). Scale bars, 1 μm. (C) Electron microscopy analysis shows that mitochondria in *Parl^-/-^* primary spermatocytes are swollen with few thin irregular cristae and loss of normal matrix density (right panel, arrowheads) compared to WT (left panel, arrowheads). The thin arrows indicate the intermitochondrial cement (nuage) typically associated with mitochondria in primary spermatocytes. The large arrows indicate fully assembled synaptonemal complexes, structures that are only detectable during the zygotene and pachytene stages of prophase I in meiosis I, (5 weeks, n=3). Scale bars, 0.5 μm.

Therefore, we examined primary spermatocytes showing fully assembled synaptonemal complexes with the typical tripartite pattern consisting of two parallel lateral regions and a central element which is only detectable during the zygotene and pachytene stages of prophase I in meiosis I^30–32^ (Fig. 2C). Compared to the mitochondria of WT primary spermatocytes, which are typically small with dilated cristae and dense finely granular matrix, mitochondria in *Parl^-/-^* spermatocytes appear consistently swollen with few thin irregular cristae and loss of normal matrix density^31^ (Fig. 2C). Abnormal mitochondrial morphology appears the earliest ultrastructural change detectable in PARL-deficient spermatocytes localized in the abluminal compartment, compared to adluminal spermatocytes where cell abnormalities also involve other membranous cell compartments including the disruption of endoplasmic reticulum, Golgi apparatus, and nuclear envelope. Chromatin clumping, and nuclear fragmentation are also evident (Fig. 2B and Fig. 2-figure supplement 1A). In sharp contrast, the ultrastructural features of other cell types in the seminiferous tubules and surrounding interstitium, including spermatogonia, Leydig, and Sertoli cells, appear normal with preserved mitochondrial morphology (Fig. 2-figure supplement 1B). Altogether, these data indicate the presence of early mitochondrial ultrastructural abnormalities culminating in extensive degeneration and death of arrested PARL-deficient spermatocytes.

### Impaired spermatogenesis in PARL-deficient testis is not driven by misprocessing of PARL substrates PINK1, PGAM5, and TTC19

Next, we asked to what extent the severe spermatogenesis defect induced by PARL deficiency can be attributed to the misprocessing and altered maturation of PARL’s substrates. To answer this question, we first tested the expression of established PARL substrates in testis. *Parl* testis mitochondria show remarkable accumulation of PINK1 and unprocessed full-length PGAM5, as well as almost total lack of the mature form of TTC19 (Fig. 3A), similar to what was previously observed in brain^10^ and in cultured cells^17^. Conversely, other PARL substrates such as DIABLO, STARD7, and CLPB show only subtle misprocessing or expression changes, likely due to compensatory proteolytic cleavage from alternative proteases (Fig. 3A). Therefore, we focused on PINK1, PGAM5, and TTC19, asking whether their genetic modulation modifies or reproduces the testicular phenotype seen in *Parl^-/-^* mice. PINK1 and PGAM5 play key functional roles in maintaining mitochondrial integrity and homeostasis and they are linked to both Parkinson’s disease and spermatogenesis defects^33–35^. TTC19 is a mitochondrial protein required for the catalytic activity of Complex III^21^ Pathogenic variants in *TTC19* cause mitochondrial disease in humans including Leigh syndrome^22^. To test this hypothesis, we analyzed testes from a series of genetically engineered mutant mouse lines incorporating multiple full gene knockouts including both *Parl* and *Pink1* (*Parl^-/-^/Pink1^-/-^*); *Parl* and *Pgam5* (*Parl^-/-^/Pgam5^-/-^*); *Pink1* and *Pgam5* (*Pink1^-/-^/Pgam5^-/-^*); *Parl, Pink1*, and *Pgam5* combined (*Parl^-/-^/Pink1^-/-^/Pgam5^-/-^*); and *Ttc19* (*Ttc19^-/-^*). As previously observed for the brain^10^, the severe testicular phenotype associated with PARL deficiency remained unmodified upon additional deletion of *Pink1* or *Pgam5* either alone or combined (Fig. 3B). On the contrary, the testes from *Pink^-/-^ Pgam5^-/-^* and *Ttc19^-/-^* appeared completely normal showing typical postpubertal spermatogenesis (Fig. 3B) and preserved reproductive activity. In conclusion, these observations indicate that impaired spermatogenesis in PARL-deficient mice is not driven by misprocessing and/or altered maturation of the substrates PINK1, PGAM5, and TTC19 despite their severely affected proteolytic processing, implicating additional pathogenetic mechanisms involved in the testicular phenotype.

**Figure 3.**
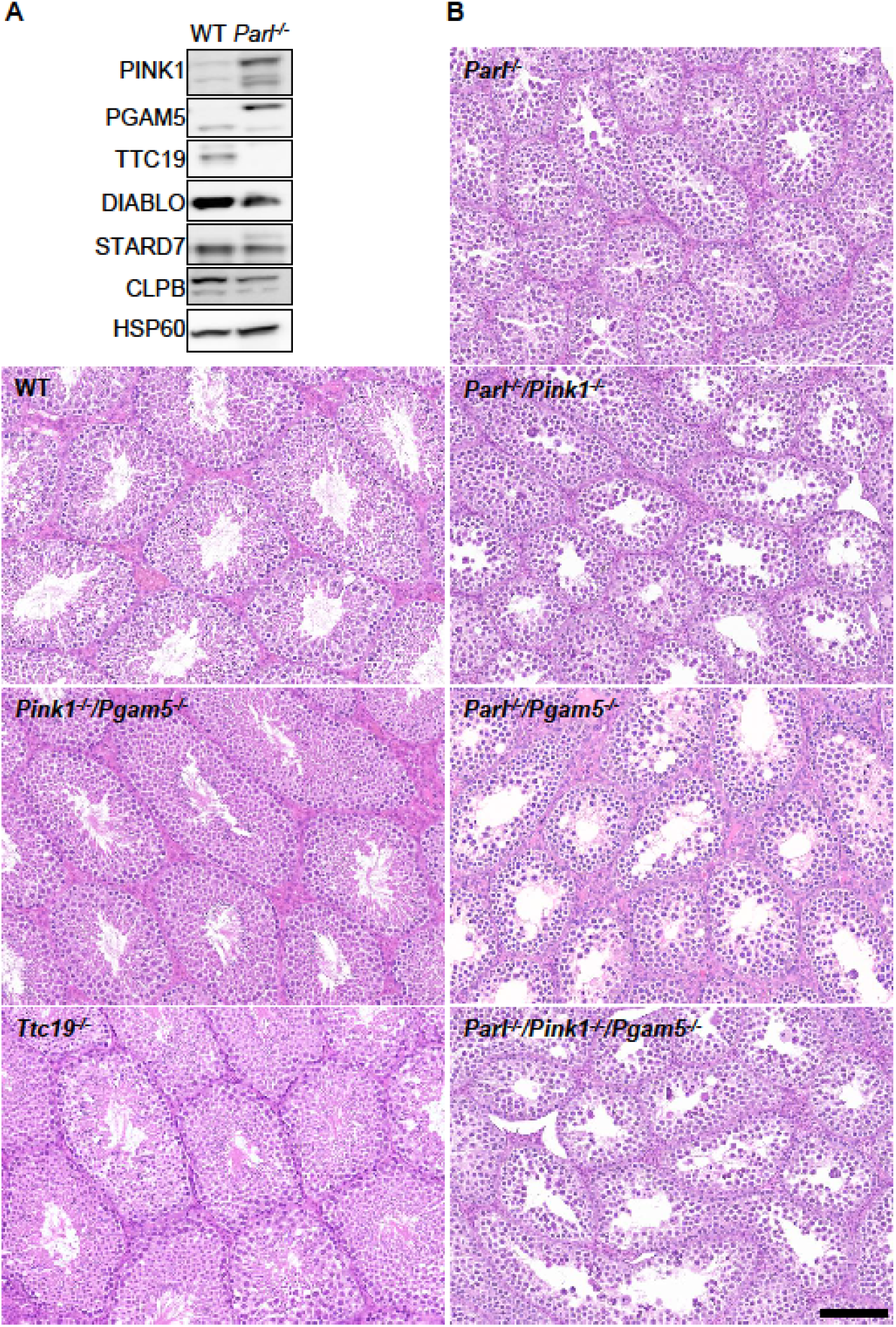
Mice with genetic manipulation of the PARL substrates PINK1, PGAM5, and TTC19 do not reproduce or modify *Parl^-/-^* testis phenotype. (A) Immunoblots of testis lysates from 6-week-old WT and *Parl^-/-^* mice with antibodies for the established PARL substrates PINK1, PGAM5, TTC19, DIABLO, STARD7, and CLPB. Severe accumulation of unprocessed PINK1 and PGAM5, as well as severe decrease in the mature processed form of TTC19 is evident in *Parl^-/-^* testis. (B) Histology of testes from 7-week-old mice of the indicated genotypes. *Parl^-/-^/Pink1^-/-^*, *Parl^-/-^/Pgam5^-/-^* and *Parl^-/-^/Pink1^-/-^/Pgam5^-/-^* show no modification of the testicular phenotype compared to *Parl^-/-^* mice. *Ttc19^-/-^* and *Pink1^-/-^/Pgam5^-/-^* mice have no evident testis pathology and are fertile (HE stain, n=3). Scale bar, 145 μm. **Figure 3—source data 1** Original images for Figure 3A.

### Mitochondria in PARL-deficient testis exhibit severe respiratory chain defects

Spermatogenesis is characterized by important metabolic adaptations and mitochondrial function plays a critical role across germ cell maturation^29^. Moreover, mitochondrial morphology and function are bidirectionally interconnected, raising the question of the functional impact of the mitochondrial morphological abnormalities identified in *Parl^-/-^* spermatocytes. Therefore, we performed a detail mitochondrial functional analysis of PARL-deficient testis mitochondria. As PARL has been previously linked to differences in mitochondrial biogenesis^36^, we wondered whether mitochondrial mass is reduced in *Parl^-/-^* testis. Expression of the outer mitochondrial membrane protein TOMM20, and of the inner membrane ATP synthase beta subunit ATPB showed similar expression levels in WT and *Parl^-/-^* testis (Fig. 4A and Fig 5B), suggesting unaltered mitochondrial mass. Similarly, mitochondrial DNA abundance was also not significantly different (Fig. 4B). Next, we asked whether mitochondrial respiratory chain complexes were normally assembled in *Parl^-/-^* testis mitochondria. Blue native gel electrophoresis shows severe assembly alterations of multiple respiratory chain complexes including Complex I, Complex III, and Complex IV, and of the supercomplex (Fig. 4C). Since respiratory chain complexes’ supramolecular assembly is required for optimizing the efficiency of mitochondrial oxidative phosphorylation, we then examined if PARL deficiency ultimately resulted in impaired mitochondrial respiration in testis mitochondria. To answer this question, we measured oxygen consumption by high-resolution respirometry in testis mitochondria supplied with substrates and specific inhibitors for Complex I (CI), Complex II (CII), and Complex IV (CIV) as illustrated in Fig. 4D. Importantly, both phosphorylating respiration, whether driven by complex I substrates only (CI OXPHOS) or by both Complex I and II (CI+II OXPHOS), and uncoupled respiration, whether driven by Complex II only (CII ET), by both Complex I and II substrates (CI+II ET), or by Complex IV (CIV), were severely diminished in *Parl^-/-^* testis mitochondria indicating a severe electron transport defect (Fig. 4E). During certain types of regulated cell death, such as apoptosis, the outer mitochondrial membrane becomes permeable, and cytochrome c is released from the mitochondrial intermembrane space to the cytosol, leading to decreased mitochondrial respiration and proteolytic activation of executioner caspases^37^. To investigate whether the mitochondrial respiratory defects of *Parl^-/-^* testis resulted from mitochondria outer membrane permeabilization, we measured the enhancement of Complex IV (CIV)-driven respiration after the addition of exogenous cytochrome c. Intact mitochondria outer membrane is impermeable to exogenous cytochrome c, whereas disruption of the outer mitochondrial membrane causes permeability to exogenous cytochrome c and consequent enhancement of CIV-driven respiration. Exogenous cytochrome c did not significantly enhance CIV-driven respiration in both WT and *Parl^-/-^* testis mitochondria (Fig. 4E), indicating lack of significant outer mitochondrial membrane permeabilization. Therefore, the identified electron transport defect is not the consequence of cytochrome c loss. Next, to obtain cell-type insights into the identified electron transport defect, we performed cytochrome c-oxidase activity staining in frozen tissue sections. We found that the function of this enzyme was significantly decreased in PARL-deficient germ cells, but not in Leydig cells (Fig. 4F). In conclusion, PARL is required to prevent severe electron transport chain defects in spermatocyte mitochondria.

**Figure 4.**
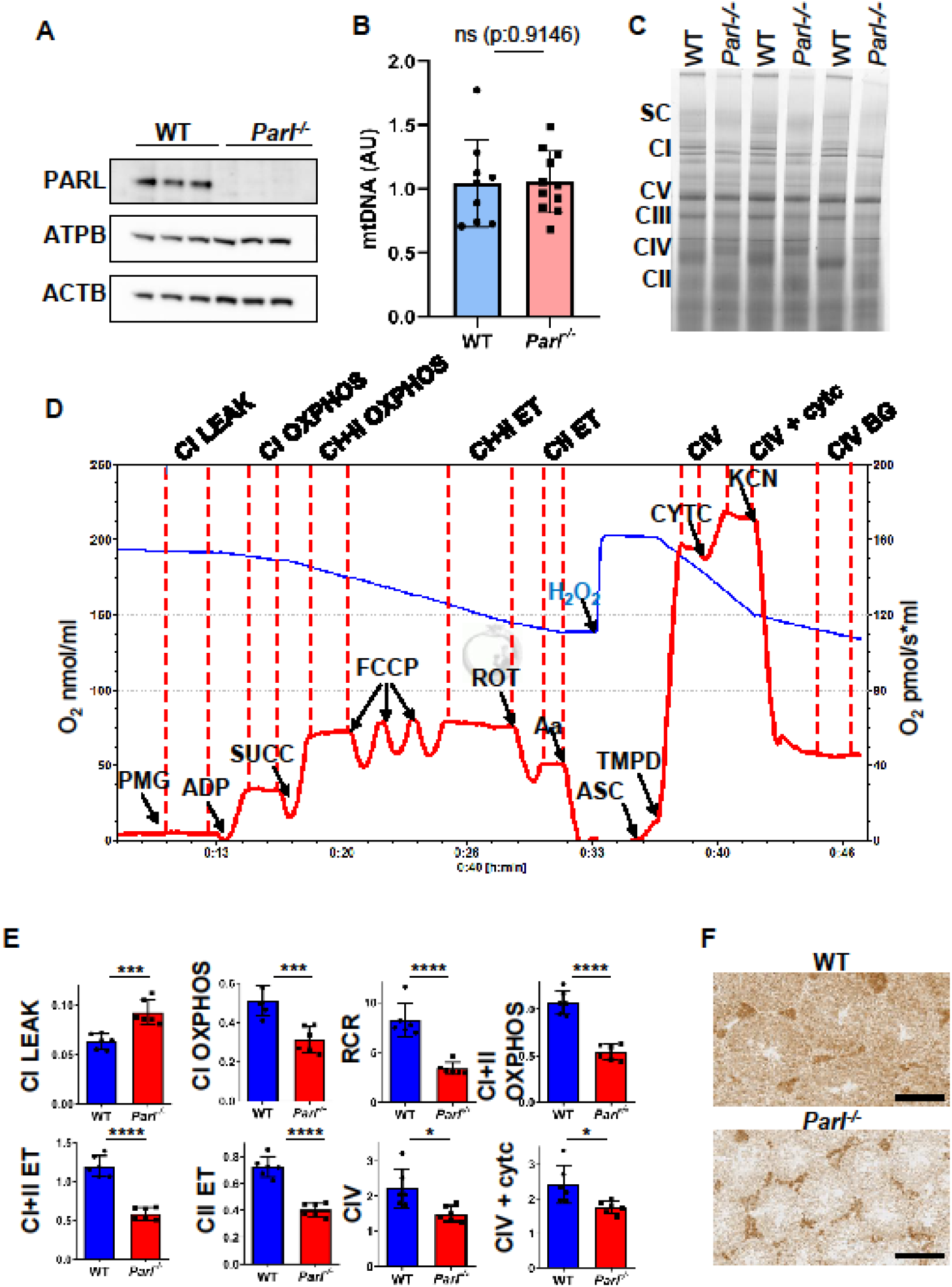
Severe mitochondrial electron transfer defects in *Parl^-/-^* testis mitochondria. (A) Immunoblots of testis lysates from 6-week-old WT and *Parl^-/-^* mice with antibodies for PARL, ATPB, TOMM20, and ACTB, (n = 3). ACTB is the loading control. (B) Quantification of mitochondrial DNA in 5-week-old WT and *Parl^-/-^* testis normalized to nuclear DNA (n=10). (C) Blue native gel electrophoresis of testis mitochondria from 6-weeks-old WT and *Parl^-/-^* mice (n=3). Mitochondrial complexes and supercomplexes are visualized after staining with Instant Blue. Assembly defects are evident for Complex I, Complex III, Complex IV, and the supercomplex. (D) Representative trace illustrating the protocol for high-resolution respirometry in testis mitochondria. The blue trace indicates the O2 concentration and the red trace indicates its time derivative. Testis mitochondria (150 μg) were loaded in Miro6 buffer. Substrates are as follows: CI (PMG, pyruvate + malate + glutamate), CII (Succ, succinate), and CIV (ASC/TMPD, ascorbate + TMPD). The uncoupler is CCCP. The specific mitochondrial inhibitors are rotenone (ROT) for Complex I, Antimycin a (Aa) for Complex III, and cyanide (KCN) for Complex IV. Respiratory states are indicated between red dashed lines. CI LEAK, CI-driven leak respiration; CI OXPHOS, CI-driven phosphorylating respiration; CI+II OXPHOS, phosphorylating respiration driven by combined activation of CI and II; CI+II ET, electron transfer capacity driven by combined CI and II; CII ET, ET driven by CII; CIV, CIV-driven respiration; Cytc, exogenous cytochrome c is added to evaluate the integrity of the outer mitochondrial membranes. H2O2 in the presence of catalase is used to reoxygenate the chamber. (E) Quantification of the respiratory states of testis mitochondria from 6-week-old WT and *Parl^-/-^* mice (n=6) as from the protocol described in (D) and in the methods section. Bar graphs indicate average ± SD. Statistical significance calculated by two-sided Student t test: *P < 0.05, ***P < 0.001, and ****P < 0.0001. (F) Cytochrome c oxidase histochemistry in frozen testis sections from 6-week-old WT and *Parl^-/-^* mice (n=3). **Figure 4—source data 1** Original images for Figure 4A.

### PARL deficiency causes impaired testicular Coenzyme Q (CoQ) biogenesis and redox

In our previous study, we showed that brain mitochondria from PARL-deficient mice have Coenzyme Q (CoQ) deficiency linked to impaired COQ4 expression, and a severe increase of the ratio between reduced and oxidized CoQ resulting from Complex III dysfunction caused by TTC19 impaired proteolysis^10^. CoQ is a hydrophobic lipid with essential cellular functions both as an electron carrier of the mitochondrial respiratory chain, and as a lipophilic free-radical-scavenging antioxidant preventing lipid peroxidation^38^. In mammalian mitochondria,

CoQ is reduced by multiple converging pathways including Complex I, Complex II, dehydroorotate dehydrogenase, sulfide-quinone oxidoreductase, and electron transfer dehydrogenase, but it is only oxidized by Complex III. CoQ is thought to play an important role in promoting testicular function and maturation of male germ cells by preventing oxidative damage^39,40^. We found that CoQ levels are significantly decreased in *Parl^-/-^* testis and the ratio between reduced and oxidized forms of CoQ (Coq red/ox) is dramatically increased compared to WT (Fig. 5A), as previously seen in brain^10^. Increased CoQ reduction most likely originates from impaired CoQH2 oxidation caused by impaired Complex III activity resulting from the severe TTC19 deficiency (Fig 3A), and Complex III assembly defects (Fig. 4E). Interestingly, as previously seen in *Parl^-/-^* brains^10^, the levels of COQ4, a protein required for coenzyme Q biogenesis^41,42^, are severely diminished in postpubertal *Parl^-/-^* testis both by WB (Fig. 5B) and IHC (Fig. 5C). COQ4 expression is diffusely decreased in different cell types of *Parl^-/-^* testis, and the deficit seems particularly prominent in *Parl^-/-^* spermatocytes, including those localized in the abluminal compartments suggesting a severe CoQ biosynthesis defect beginning in the early stages of degeneration (Fig. 5C). Decreased COQ4 expression is also evident in Leydig and Sertoli cells (Fig. 5C). Altogether, our data indicate that PARL is required for the normal expression of COQ4, essential to maintain CoQ biosynthesis and for maintaining a balanced CoQ red/ox ratio.

**Figure 5.**
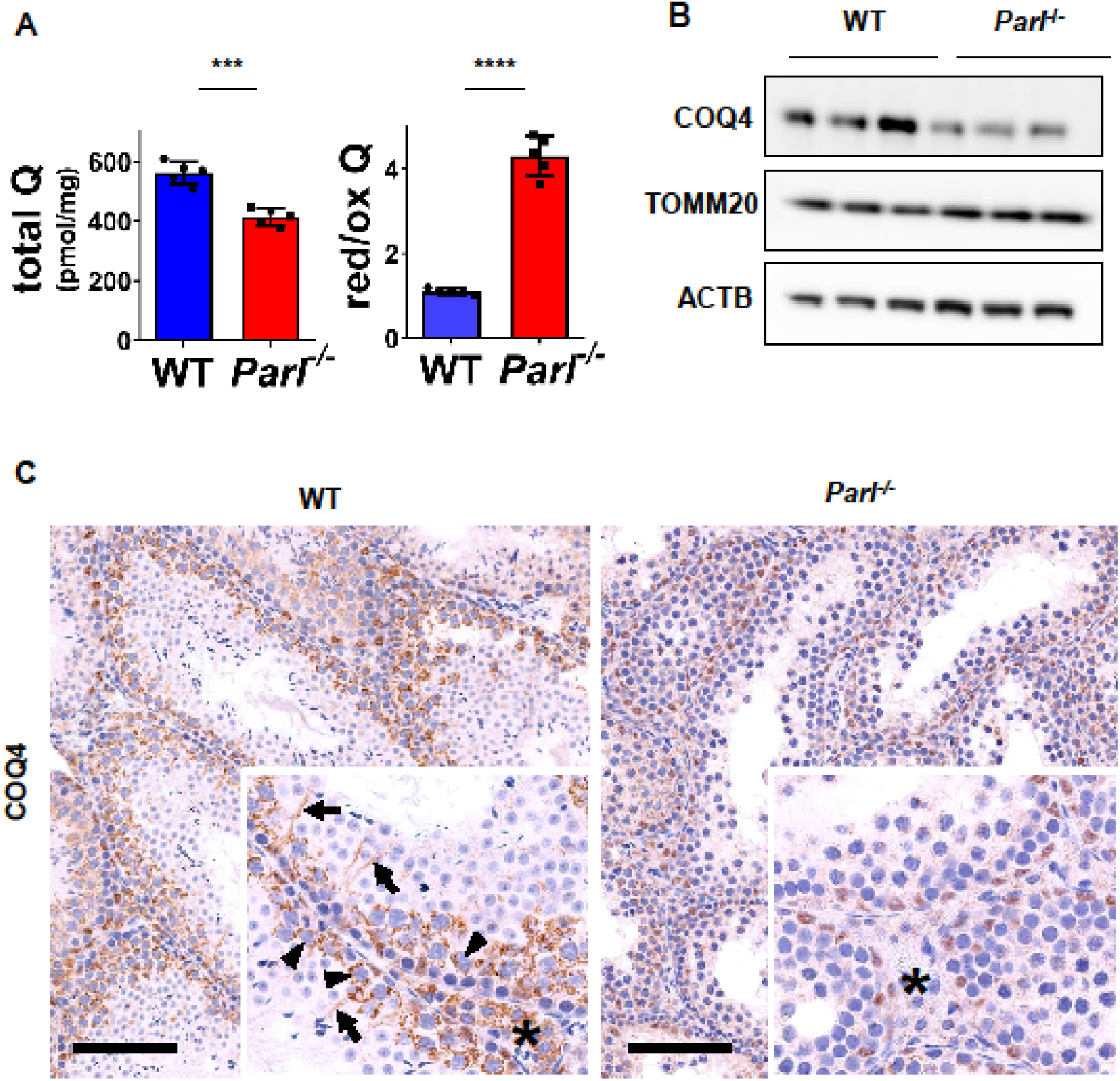
Severe alteration in Coenzyme Q biosynthesis and redox state in *Parl^-/-^* testis. (A) Concentration (left) and CoQ red/ox ratio (right) of total CoQ (Q_9_+Q_10_) measured by HPLC in the testes of 5-week-old WT and *Parl^-/-^* mice (n=5). Total CoQ levels are severely decreased in *Parl^-/-^* testis compared to WT. Moreover, the redox status is altered with drastic elevation in the reduced form of CoQ. Bar graphs indicate average ± SD. Statistical significance calculated by two-sided Student *t*-test: ***P < 0.001, and ****P < 0.0001. (B) Immunoblot analysis of testis total lysates obtained from 5 weeks-old WT and *Parl^-/-^* mice with antibodies for COQ4, TOMM20, and ACTB (n=3). ACTB is the loading control. Decreased COQ4 expression is evident. (C) Immunohistochemistry for COQ4 shows severely reduced levels of COQ4 expression in 6-week-old *Parl^-/-^* testis as compared to WT controls (n=3). The deficit particularly prominent in *Parl^-/-^* arrested spermatocytes, almost devoid of COQ4 expression, as compared to the high constitutive levels of COQ4 expression observed in WT spermatocytes (inset, stage II tubule, arrowheads). Decreased COQ4 expression is also evident in *Parl^-/-^* Leydig cells as compared to WT mice (insets, asterisk). In addition, COQ4-positive Sertoli cell projections observed in WT mice (inset, stage II tubule, arrows) are not evident in the seminiferous tubules of *Parl^-/-^* mice. Scale bar, 100 μm. **Figure 5—source data 1** Original images for Figure 5B.

### PARL deficiency leads to ferroptosis in arrested spermatocytes

Next, we wondered which specific cell death modality was responsible for the severe germ cell degeneration and demise observed in PARL-deficient mice. The prominent features of chromatin condensation and nuclear fragmentation in adluminal germ cells during the late stages of degeneration (Fig. 2A and 2B), and previous links of PARL to antiapoptotic properties *in vitro*^23^, lead us to assess the potential involvement of increased apoptosis in this phenotype. However, levels of caspase-3 activation in the seminiferous tubules of *Parl^-/-^* mice is comparable to WT, failing to provide evidence for the implication of apoptosis (Fig. 2-figure supplement 2). The identification of decreased CoQ concentration (Fig. 5A-C) and the presence of severe ultrastructural abnormalities involving mitochondria and other membranous cell compartments (i.e., ER, Golgi apparatus, and nuclear envelope) (Fig. 2C and Fig. 2-figure supplement 1A), led us to hypothesize the possible implication of ferroptosis, a programmed cell death modality characterized by lipid peroxidation of cell membranes^43,44^. Previous studies conducted in cultured cells have in fact demonstrated the importance of CoQ producing mevalonate pathway^45^ and of CoQ reducing pathways driven by FSP1^46,47^, DHODH^48^, and GCH1^49^ in this programmed cell death modality. To test this hypothesis, we checked the expression of GPX4, an essential antioxidant peroxidase, required for preventing ferroptosis by directly reducing phospholipid hydroperoxide in cell membranes using reduced glutathione as substrate^50,51^. Notably, immunoblot analysis shows that GPX4 expression is almost completely abolished in *Parl^-/-^* testis (Fig 6A). Next, to investigate the effect of PARL proteolytic activity on GPX4 expression, we checked GPX4 expression in mouse embryonic fibroblasts with and without catalytically active or inactive PARL. The results do not show modifications of GPX4 migration or expression and therefore do not indicate GPX4 as a direct substrate of PARL (Figure 6-figure supplement 1A). Moreover, the striking effect of PARL deficiency on GPX4 expression does not appear to be generalized in other organs (Fig. 6-figure supplement 1B). Using IHC to gain cell-type specific insights, we show that GPX4 expression is specifically abrogated in *Parl^-/-^* spermatocytes but not Leydig cells (Fig. 6C, top panels). Next, we established that other known mediators of ferroptosis, such as cellular tumor antigen p53 (also referred to as p53), a master regulator of both canonical and non-canonical ferroptosis pathways^52,53^, and transferrin receptor protein 1 (TfR1), which mediates cellular uptake of iron via receptor-mediated endocytosis^54^, were significantly upregulated (Fig. 6C and Fig. 6-figure supplement 2). Using IHC, we detected prominent p53 nuclear expression mainly in adluminal and exfoliated degenerating spermatocytes from PARL-deficient testis, while testicular p53 levels are constitutively very low or undetectable in normal postpubertal mice^55^ (Fig. 6-figure supplement 2). In WT testis, expression of TfR1, is normally very high in spermatogonia and drastically decreases in spermatocytes and subsequent maturation forms^56,57^ (Fig. 6C, middle panel). Conversely, *Parl^-/-^* mice show persistent overexpression of TfR1in arrested spermatocytes suggesting abnormally high iron uptake (Fig. 6C, middle panels). TfR1 overexpression is particularly prominent in adluminal and exfoliated spermatocytes during the late stages of degeneration (Fig. 6C, middle panels). We further investigated whether the observed severe reduction in GPX4 and CoQ levels ultimately resulted in increased lipid peroxidation, the ultimate biochemical outcome of ferroptosis. Immunoblot with an antibody specific for 4-hydroxynonenal (HNE) adducts, the most abundant and stable end-products of lipid peroxidation, show significantly increased HNE signal in *Parl^-/-^* testis (Fig. 6B), but not in brain (Fig. 6-figure supplement 1C), compared to WT confirming a testis specific effect of PARL deficiency on lipid peroxidation. Interestingly, IHC analysis shows that the accumulation of HNE gradually increases from the abluminal to the adluminal and exfoliated *Parl^-/-^* spermatocytes (Fig. 6C, bottom panels) reflecting the same distribution and progression described above for TfR1 and p53 overexpression. Taken together, these findings indicate that ferroptosis is a cell-type-specific downstream effect of PARL deficiency and the mechanism responsible for the degeneration and demise of arrested spermatocytes in *Parl^-/-^* testis.

**Figure 6.**
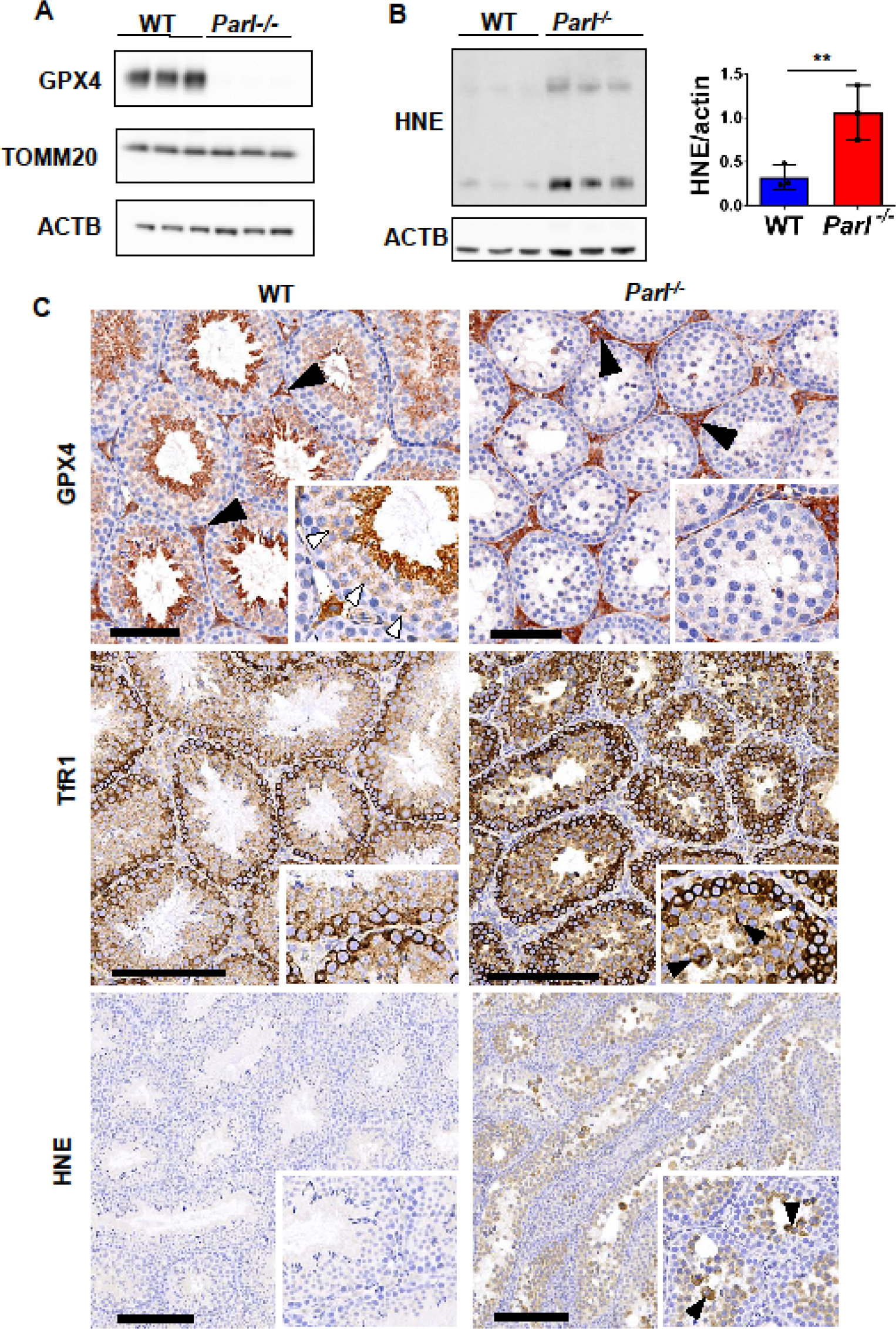
Massive ferroptosis activation in *Parl^-/-^* arrested spermatocytes. (A) Immunoblot of testis total lysates obtained from 5-weeks-old WT and *Parl^-/-^* mice using antibodies for GPX4, TOMM20, and ACTB (n=3). ACTB is the loading control. GPX4 expression is barely detectable in *Parl^-/-^* testis. (B) Immunoblot analysis of total testis lysates from 7 weeks-old WT and *Parl^-/-^* mice using anti-HNE and anti-ACTB antibodies (n=3). ACTB is the loading control. Quantification of the HNE/ACTB ratio is shown on the right as graphs indicating average ± SD (n=3, **P < 0.01). The statistical significance HNE/ACTB ratio increase in *Parl^-/-^* mice has been calculated by two-sided Student *t*-test. (C) Immunohistochemistry for GPX4, TfR1, and HNE in testis from 6-week-old WT and *Parl^-/-^* mice (n=3). GPX4 expression is barely detectable in *Parl^-/-^* arrested spermatocytes compared to WT (inset, stage X tubule, white arrowheads), while it is unaffected in interstitial Leydig cells (black arrowheads) (top panels, scale bar, 100 um). In WT testis, TfR1 expression drastically decreases as basal spermatogonia mature into spermatocytes; in contrast, TfR1 expression in *Parl^-/-^* testis is abnormally increased in arrested spermatocytes (middle panels; scale bars, 200 μm), especially in adluminal/exfoliated degenerating cells (inset, arrowheads). A similar pattern is also observed for HNE immunohistochemistry, which shows gradual intensification of lipid peroxidation during spermatocyte degeneration with prominent signal in adluminal/exfoliated spermatocytes (inset, arrowheads) (bottom panels; scale bar, 200 μm). **Figure 6—source data 1** Original images for Figure 6A. **Figure 6—source data 2** Original images for Figure 6B.

## DISCUSSION

This work reveals an essential role of PARL in maintaining spermatogenesis and germ cell survival. The testicular phenotype represents the earliest manifestation of PARL deficiency, which specifically affects primary spermatocytes leading to a maturation arrest before the completion of the first meiotic division. Interestingly, male infertility and spermatogenesis defects had been reported in Drosophila mutant for the mitochondrial rhomboid orthologue Rhomboid-7^58^ indicating that this phenotype is highly conserved across different Phyla. In the mouse, the mechanisms leading to the testicular phenotype appears to involve severe mitochondrial functional and ultrastructural defects involving marked defects at the level of the mitochondrial electron transport chain and associated with localized induction of ferroptosis in arrested spermatocytes. These results are in line with and extend our previous observations in PARL-deficient brain mitochondria reporting defective Complex III activity and altered CoQ biosynthesis caused by impaired COQ4 expression. In the testis the respiratory chain defects are even more severe than in brain and include multiple assembly abnormalities of complex I, III, and IV, that were not present in the nervous system^10^. In both testis and brain these defects are associated with an increased proportion of reduced versus oxidized CoQ. Within testis, the respiratory chain defect and the COQ4 expression deficit appear particularly severe in primary spermatocytes compared to other cell types. Mitochondrial morphological abnormalities appear also restricted to spermatocytes. Altogether, these data confirm a crucial role of PARL in the maintenance of the respiratory chain, mitochondrial ultrastructure, and CoQ biosynthesis^10^. We believe that this severe respiratory chain defect represents the driving mechanism of the observed meiotic arrest, by making primary spermatocytes unable to accommodate the sudden bioenergetic shift, from glycolytic to oxidative, required during the first meiotic division to cope with the increased energy demand^59^. This hypothesis is consistent with reported spermatogenesis defects in other mouse models characterized by different types of mitochondrial insults, including defective mitochondrial DNA^60,61^, adenylates transport^62^, cardiolipin biosynthesis^63^, mitochondrial dynamics^64,65^, and mitochondrial proteolysis^66,67^. Altogether mitochondrial fitness appears crucial for ensuring germ cell differentiation during spermatogenesis.

An essential role of PARL in cell survival has been established since its original description, because of its lethal phenotype in germline knockout mice^23^ and other organisms^14^, with contradictory links to apoptosis^17,23^ in cellular models. More recently we reported that PARL deficiency does not affect apoptosis in the brain, but it induces necrosis. Similarly, apoptosis is not implicated in the testicular phenotype of PARL-deficient mice. Importantly, our study establishes the specific induction of ferroptosis in PARL-deficient spermatocytes that is responsible for their final demise. Ferroptosis represents a specific form of regulated cell death that is characterized by uncontrolled iron-dependent lipid peroxidation of cell membranes^49,50^. It can be experimentally induced in vitro by inhibition of the phospholipid-hydroperoxide glutathione peroxidase GPX4, the master regulator of ferroptosis, or depletion of its substrate glutathione^56,57^. GPX4 exists in three distinct isoforms originating from different transcription initiation sites: a full-length mitochondrial form, a shorter cytosolic form, and a nuclear isoform^68^. GPX4 expression is highest in testis, where the predominant form is represented by the mitochondrial isoform^69^. The crucial physiological relevance of GPX4 is demonstrated by the observation of cell death in mice lacking GPX4. Germline deletion of *Gpx4* is embryonically lethal^70^. Conditional deletion of *Gpx4* in different tissues, such as the nervous system^71^, kidney^72^, liver^73^, and endothelium^74^ show severe pathological phenotypes. Importantly, spermatocyte-specific deletion of *Gpx4* in mice leads to severe testicular atrophy, reduced spermatogenesis, germ cell death, and infertility^75^, indicating that GPX4 plays an important role in male reproductive biology. In addition, GPX4 activity is severely reduced in the sperm of infertile patients highlighting an important role of GPX4 in human spermatogenesis^76,77^. An additional GPX4 independent anti-ferroptotic pathway relies on CoQ that provides powerful protection from lipid peroxidation through its reduced form^45,47–49,78^. CoQ is most abundant in mitochondria, where it is synthetized, but it is also present in other cell membranes including the plasma membrane, Golgi, and endoplasmic reticulum^42,79^. The contribution of mitochondria to ferroptosis is still debated^80^, but recent evidence, including the data here provided, indicates that mitochondria play important roles in this process^81^. A negligible role of mitochondria in ferroptosis has initially been argued based on *in vitro* observations, since ferroptosis can be induced in cultured cells devoid of mitochondrial DNA^82^ or artificially deprived of mitochondria through overexpression of PARKIN and addition of large doses of mitochondrial uncouplers^83^. Another study proposed a promoting role of the mitochondrial respiratory chain based on the suppression of ferroptosis induced by cysteine deprivation in mouse embryonic fibroblasts through the addition of inhibitors targeting any of the respiratory chain complex I, II, III, or IV^84^. Conversely, in cancer cells treated with GPX4 inhibitors to induce ferroptosis, dihydroorotate dehydrogenase DHODH, an inner membrane enzyme involved in pyrimidine biosynthesis, has been recently found to suppress ferroptosis by reducing CoQ^48^ suggesting that mitochondrial pathways can modulate ferroptosis. Although much of what is known today about ferroptosis comes from *in vitro* experiments^48^, its pathophysiological implication in diseases is emerging^85^. In hearts from mice with different types of mitochondrial dysfunction, such as mitochondrial genome expression defects ^86^ or cytochrome c oxidase deficiency^81^, GPX4 expression has been shown to increase. In a recent study, Ahola and collaborators have elegantly shown that upregulation of GPX4 provides a crucial homeostatic response to prevent ferroptosis in heart tissue affected by OXPHOS deficiency^81^. This adaptation is part of a broad response to mitochondrial dysfunction called integrated stress response, which is mediated by the transcription factor ATF4 and involve induction of the trans-sulphuration pathway to promote glutathione metabolism and increased incorporation of selenium supporting increased GPX4 expression ^81^. Impairing this homeostatic response to OXPHOS deficiency by knocking out either the mitochondrial protease OMA1 or its substrate DELE1 aggravated cardiomyopathy induced by COX10 deficiency by decreasing GPX4 to basal levels, comparable to WT, and inducing ferroptosis^81^. These data clearly demonstrate the physiological importance of homeostatic mechanisms to prevent ferroptosis in conditions of defective oxidative phosphorylation. In our model, we describe a different response, where spermatocytes affected by OXPHOS deficiency induced by *Parl* ablation are unable to express GPX4. The absence of PARL activates ferroptosis in spermatocytes through severe simultaneous effects on GPX4 and CoQ, the two major and independent ferroptosis regulatory pathways. Interestingly, some level of interdependence between these two pathways is suggested by similar simultaneous inhibitory effects of the ferroptosis inducer FIN56^45^ and by influence of the mevalonate pathways on the isopentenylation of selenocysteine-tRNA^87^ that is required for efficient GPX4 expression. The dramatic spermatocytes degeneration that we report is consistent with the lethality of GPX4 deficiency in the context of respiratory chain deficiency, which has previously been demonstrated in genome wide CRISPR screens in cultured cells treated with a broad spectrum of mitochondrial respiratory chain inhibitors^88^. Interestingly, GPX4 is not a PARL substrate, and its absence is not the simple result of impaired proteolysis. The reason why only spermatocytes undergo ferroptosis in absence of PARL seems related to the restricted deficiency of GPX4 in this cell type. The distinct vulnerability of spermatocytes is also likely influenced by their severe CoQ biosynthesis defect, and their particularly high poly-unsaturated fatty acid content^89^, which might make them exceptionally susceptible to lipid peroxidation.

Altogether, these data provide evidence that ferroptosis can be initiated by a primary mitochondrial defect *in vivo*. This may have important therapeutical implications in the future as effective ferroptosis inhibitors *in vivo* will be available. Moreover, this study illustrates how specific phenotypes resulting from systemic mitochondrial dysfunctions can be caused by cell-type specific downstream pathophysiological mechanisms. This is important to improve our understanding of how differential tissue susceptibility shapes the different manifestations of mitochondrial diseases as well as their potential treatments. In conclusion, this work reveals a crucial role of PARL in spermatogenesis by shaping the mitochondrial electron transport chain, mitochondrial ultrastructure, CoQ biosynthesis, and GPX4 expression in spermatocytes which is required for prevention of ferroptosis. Importantly, this study prompts further investigations on ferroptosis as an emerging pathomechanism and potential pharmacological target of mitochondrial diseases and other disorders causing male infertility.

## METHODS

### Animals and Husbandry

Mice with full knockout germline deletion of *Parl* (*Parl^-/-^*) (MGI:3693645), *Pgam5* (*Pgam5^-/-^*) (MGI:5882561), *Pink1* (*Pink1^-/-^*) (MGI:5436308), *Ttc19* (*Ttc19^-/-^*) (MGI:6276545), and conditional *Parl* ablation under the nestin promoter (*Parl^L/L^::Nes^Cre^*) (MGI:3526574, MGI:2176173) have been generated as previously described^10,23^. All mutant mouse lines were maintained on a C57BL/6J background. Mice were kept in a SPF facility and multiply housed in filter top polycarbonated cages enriched with wood-wool and shavings as bedding. Standard rodent diet and acidified tap water were provided *ad libitum*. Animal rooms were maintained at 22°C ± 2°C with a 45% and 70% relative humidity range, 50 air changes per hour, and twelve-hour light/dark cycles. Mice were included in a health-monitoring program developed in accordance with guidelines of the Federation of European Laboratory Animal Science Associations (FELASA). All experiments were approved by the Ethical Committee on Animal Experimenting of the University of Leuven (IACUC protocol #072/2015) and by the French Ministry (DUO-OGM 5769 29/3/2019).

### Pathological and immunohistochemical examination

Testes harvested from postpubertal mutant mice and WT matched controls were immersion-fixed in 10% neutral buffered formalin for 24-48 hours at room temperature (RT). Samples were then routinely processed for paraffin embedding, sectioned at 5 μm, and stained with hematoxylin and eosin (HE) for histopathological assessment. For immunohistochemistry (IHC), 5 μm thick paraffin sections were mounted on ProbeOn™ slides (Thermo Fisher Scientific #15-188-51)). The immunostaining procedure was performed using a Leica BOND RXm automated platform combined with the Bond Polymer Refine Detection kit (Leica #DS9800). Briefly, after dewaxing and rehydration, sections were pretreated with the epitope retrieval BOND ER2 high pH buffer (Leica #AR9640) for 20 minutes at 98°C. Endogenous peroxidase was inactivated with 3% H_2_O_2_ for 10 minutes at RT. Nonspecific tissue-antibody interactions were blocked with Leica PowerVision IHC/ISH Super Blocking solution (PV6122) for 30 minutes at RT. The same blocking solution also served as diluent for the primary antibodies. Rabbit primary antibodies against Synaptonemal Complex Protein 1 (SCP-1, Abcam ab175191, working concentration 1/200), Allograft Inflammatory Factor 1 (AIF-1, Wako 019-19741, RRID:AB_839504, working concentration 1/1200), Coenzyme Q4 (COQ4, Proteintech 16654-1AP, RRID:AB_2878296, working concentration 1/200), Glutathione Peroxidase 4 (GPX4, Sigma HPA047224, RRID:AB_2679990, working concentration 1/100), 4-Hydroxynonenal (HNE, Alpha Diagnostic International HNE11-S, RRID:AB_2629282, working concentration 1/3000), transferrin receptor protein 1 (TfR1, Abcam ab214039, RRID:AB_2904534, working concentration 1/1000), Tumor Antigen p53 (p53, Leica/Novocastra NCL-L-p53-CM5p, RRID:AB_2895247, working concentration 1/300) were incubated on the sections for 45 minutes at RT. A biotin-free polymeric IHC detection system consisting of HRP conjugated anti-rabbit IgG was then applied for 25 minutes at RT. Immunoreactivity was revealed with the diaminobenzidine (DAB) chromogen reaction. Slides were finally counterstained in hematoxylin, dehydrated in an ethanol series, cleared in xylene, and permanently mounted with a resinous mounting medium (Thermo Scientific ClearVueTM coverslipper). Negative controls were obtained by replacement of the primary antibodies with irrelevant isotype-matched rabbit antibodies. HE and IHC-stained slides were evaluated by two board-certified veterinary pathologists (ER and CAA) with extensive expertise in mouse pathology. Staging of the seminiferous tubules was performed according to well-established morphological criteria^90,91^. The Aperio Versa 200 instrument was used for image acquisition.

### Immunoblot analysis

Testis total lysates were prepared by homogenization with a glass-to-glass potter homogenizer on ice in 20 mM HEPES, 100 NaCl, pH 7.4, supplemented with protease and phosphate inhibitors (ROCHE). The lysate was then transferred to a fresh tube, supplemented with Triton-X 1%, SDS 0.1%, and passed several times through a 26-gauge syringe. The samples were then centrifuged at 20,000 g for 15 minutes at 4°C to remove insoluble material. Tissue extracts or enriched mitochondrial membranes were separated in reducing and denaturing conditions in NuPage gels (Invitrogen). Proteins were transferred to PVDF 0.45 μm membranes, blocked with milk 5% TRIS-buffered saline, Tween-20 0,1% (TTBS), and incubated with the indicated primary antibodies, washed in TTBS incubated for 1 hour at room temperature with horseradish peroxidase conjugated secondary antibodies in 5% milk-TTBS or Alexa Fluor conjugated secondary antibodies. Proteins were identified by chemiluminescence or by fluorescence according to the type of secondary antibody. A PARL carboxy-terminal antibody was generated in house as previously reported^23^ (PMID: 16839884). In addition, the following commercial available antibodies were employed: anti-ACTB (Sigma A5441, RRID:AB_476744), anti-HSPD1 (HSP60) (BD Biosciences 611562, RRID:AB_399008), anti-SDH (Abcam ab14715, RRID:AB_301433), anti-ATP5B (ATP synthase-beta) (Abcam ab14730, RRID:AB_301438), anti-TOMM20 (Santa Cruz sc-11415, RRID:AB_2207533), anti-PINK1 (Cayman 10006283, RRID:AB_10098326), anti-PGAM5 (Sigma HPA036979, RRID:AB_10960559), anti-TTC19 (Sigma HPA052380, RRID:AB_2681806), anti-COQ4 (Proteintech 16654-1-AP, RRID:AB_2878296), anti-CLPB (Proteintech 15743-1-AP, RRID:AB_2847900), anti-STARD7 (Proteintech 15689-1-AP, RRID:AB_2197820), anti-DIABLO (Cell Signaling Technology 15108, RRID:AB_2798711), anti GPX4 (R&D systems MAB5457, RRID:AB_2232542;Santa Cruz sc-166570, RRID:AB_2112427), anti HNE (R&D system 198960, RRID:AB_664165), anti-Citrate synthase (Abcam ab96600, RRID:AB_10678258).

### Subcellular fractionation methods

To prepare testis enriched mitochondrial fractions for western blotting or blue native gel electrophoresis, freshly collected testis was homogenized with a motor-driven Teflon pestle set at 800 rpm in a glass potter containing ice-cold 20 mM HEPES, 225 mM sucrose, 75 mM mannitol, 1 mM EGTA pH 7.4, on ice. For mitochondrial respiration experiments, fresh testis was homogenized manually with a Teflon pestle in ice-cold 20 mM HEPES, 225 mM sucrose, 75 mM mannitol, 1 mM EGTA pH 7.4, on ice, then gently passed 5 times through a 22-gauge syringe. The homogenate was centrifuged at 700 g for 10 minutes at 4°C to remove nuclei and unbroken debris. The supernatant (tissue homogenate) was then centrifuged at 10’000 g for 10 minutes at 4°C to pellet mitochondrial enriched mitochondrial membranes. To prepare liver enriched mitochondrial fractions, freshly collected liver was thoroughly rinsed in homogenization buffer, then homogenized with a motor-driven Teflon pestle set at 800 rpm in a glass potter containing ice-cold 20 mM HEPES, 225 mM sucrose, 75 mM mannitol, 1 mM EGTA pH 7.4, on ice. The homogenate was centrifuged at 1000 g for 10 minutes at 4°C to remove nuclei and unbroken debris. The supernatant (tissue homogenate) was then centrifuged at 6,000 g for 10 minutes at 4°C. Brain mitochondria were purified according to Sims’ method^92^.

### Blue native gel electrophoresis

Blue-native-gel electrophoresis of digitonin-solubilized mitochondria was performed as described^93^. 100 μg isolated mitochondria were solubilized with 600 μg digitonin in Invitrogen Native Page sample buffer on ice for 20 minutes, then centrifuged at 20,000 g for 20 minutes at 4°C. 0,75% Coomassie G-250 was added to supernatants, which were loaded on a 3-12% gradient Invitrogen Native Page gel according to the instructions. After electrophoresis, mitochondrial complexes and super complexes were visualized by protein staining with InstantBlue^®^ Coomassie Protein Stain (ISB1L) (Abcam ab119211).

### High-resolution respirometry

Mitochondrial respiration in testis mitochondria respiration was measured in Miro6 Buffer ^94^ (20 mM HEPES, 110 mM sucrose, 10 mM KH2PO4, 20 mM taurine, 60 mM lactobionic acid, 3 mM MgCl2, 0.5 EGTA, pH 7.1, 1 mg/ml fatty acid free BSA, catalase 280 U/ml) at 37°C as previously described^10^ (PMID: 30578322). When needed H202 was added to reoxygenate the chambers by catalase mediated O2 generation. 150 μg of mitochondrial enriched membranes were loaded into the Oroboros 2K oxygraph. A typical experiment is illustrated in Fig.4D. Oxygen consumption rates were measured before and after addition of the following sequence of substrates and specific inhibitors: 1) 2.5 mM pyruvate, 10 mM glutamate, and 1 mM malate to measure Complex I-driven leak respiration (CI leak); 2) 2.5 mM ADP to determine complex I-driven phosphorylating respiration (CI OXPHOS). 3) 5 mM succinate to determine the phosphorylating respiration driven by simultaneous activation of complex I and II (CI+II OXPHOS); 4) Titrating concentrations of the mitochondrial uncoupler CCCP to reach the maximal uncoupled respiration (CI+II electron transfer capacity, ET); 5) 200 nM rotenone to fully inhibit complex I-driven respiration and measure complex II-driven uncoupled respiration (CII electron transfer capacity, CII ET); 6) 0.5 μM Antimycin A to block mitochondrial respiration at the level of complex III. Residual oxygen consumption was always negligible. 7); 2 mM ascorbate, 0.5 mM TMPD to measure cytochrome c oxidase (CIV)-driven respiration; 8) 125 μg/ml cytochrome c to evaluate mitochondrial outer membrane integrity 9) 500 μM potassium cyanide (KCN) to specifically block cytochrome c oxidase activity and measure residual background oxygen consumption caused by chemical reaction between ascorbate and TMPD. Cytochrome c oxidase-driven respiration was calculated as the cyanide sensitive oxygen consumption.

### CoQ analysis

CoQ content and the ratio of the reduced vs. oxidized forms were measured as previously described^95^.

### mtDNA copy number quantification

For mtDNA quantification, total DNA was isolated from 20-30 mg of testis tissues by using a DNeasy Blood and tissues kit (Qiagen). qPCRs were performed in triplicate in 96-well reaction plates (Applied Biosystems). Each reaction (final volume 10 μl) contained 25 ng DNA, 5 μl of Power SYBR-Green PCR Master Mix (Applied Biosystems) and 0.5 μM of each forward and reverse primer. COX1, mitochondrial encoded gene, was amplified and β2 microglobulin (β2 m), nuclear encoded gene, was used as a normalizing control. Fold changes in mtDNA amount were calculated with the ΔΔCt method. The employed primers sequences were Cox1-Mus-F: TTTTCAGGCTTCACCCTAGATGA, Cox1-Mus-R: CCTACGAATATGATGGCGAAGTG, B2M-Mus-F: ATGGGAAGCCGAACATACTG, B2M-Mus-R:CAGTCTCAGTGGGGGTGAAT

### Electron microscopy

Testis of the indicated genotype were collected and immediately fixed with 2.5% glutaraldehyde, 2% paraformaldehyde in 0.1 M cacodylate buffer pH 7.4. Tissue was stored overnight at 4°C in the fixative solution, washed in 0.1 M cacodylate buffer and post-fixed for 2 hours at RT with 1% OsO4, 1.5% K_4_Fe(CN)_6_ in 0.1 M cacodylate buffer. Sections were rinsed, stained with 3% uranyl acetate for 1 hour at 4°C and dehydrated in graded ethanol concentrations and propyleneoxide, followed by embedding in Epon^®^ Resin. Resin blocks were sectioned on a ultramicrotome. Post-staining was performed with 3% uranyl acetate followed by lead citrate staining. Semithin sections were collected on slides and stained with 1% Toluidine blue solution (Sigma-Aldrich). Ultrathin sections (60 nm) were mounted on copper grids and imaged using a JEOL transmission electron microscope.

### Cultured cells

Immortalized mouse embryonic fibroblasts (MEFs) derived from WT and *Parl^-/-^* male mice were cultured in Dulbecco’s modified Eagle’s medium/F-12 (Gibco) containing 10% fetal bovine serum (Gibco). At 30–40% confluence, the MEFs were transduced using a replication defective recombinant retroviral expression system (Clontech) with either wild-type (*Parl* WT) or with catalytic inactive *Parl S275A* as previously described^10^. Cell lines stably expressing the desired proteins were selected based on their acquired resistance to 5 μg/ml puromycin.

### Statistical analysis

Numerical data are expressed as mean ± SD from biological replicates. No statistical tests were used to predetermine sample size. Replicates numbers were decided from experience of the techniques performed and practical considerations. Two-sided student’s t test was used to compare differences between two groups using GraphPad. Differences were considered statistically significant for p ≤ 0.05. No data were excluded.

## ACKNOWLEDGMENTS

This study was supported by the University of Pennsylvania URF research funding to ER (URF Fall 19-0914) and AFM Telethon to MS (23019); the authors affiliated with the Penn Vet Comparative Pathology Core are partially subsidized by the Abramson Cancer Center Support Grant (P30 CA016520); the Aperio Versa 200 scanner used for imaging was acquired through an NIH Shared Instrumentation Grant (S10 OD023465-01A1); we are profoundly grateful to Prof. Bart De Strooper, KU Leuven, for his support and for the generous gift of all mouse strains used in this project; we thank Prof. Jeremy Wang, University of Pennsylvania, for his insightful comments.

## AUTHORS CONTRIBUTIONS

Conceptualization, E.R. and M.S.; Methodology, E.R., M.S.; Investigation, E.R., C.A.A., E.B, F.M., S.K., A.G., GL.L., P.N., and M.S.; Formal Analysis, E.R., C.A.A., and M.S.; Resources, E.R. and M.S.; Writing – Original Draft, E.R. and M.S.; Writing – Review & Editing, E.R., G.L., P.N., and M.S.; Visualization: E.R. and M.S.; Supervision, E.R. and M.S.; Project Administration, E.R. and M.S.; Funding Acquisition, E.R. and M.S.

---

**Figure 1-figure supplement 1.**
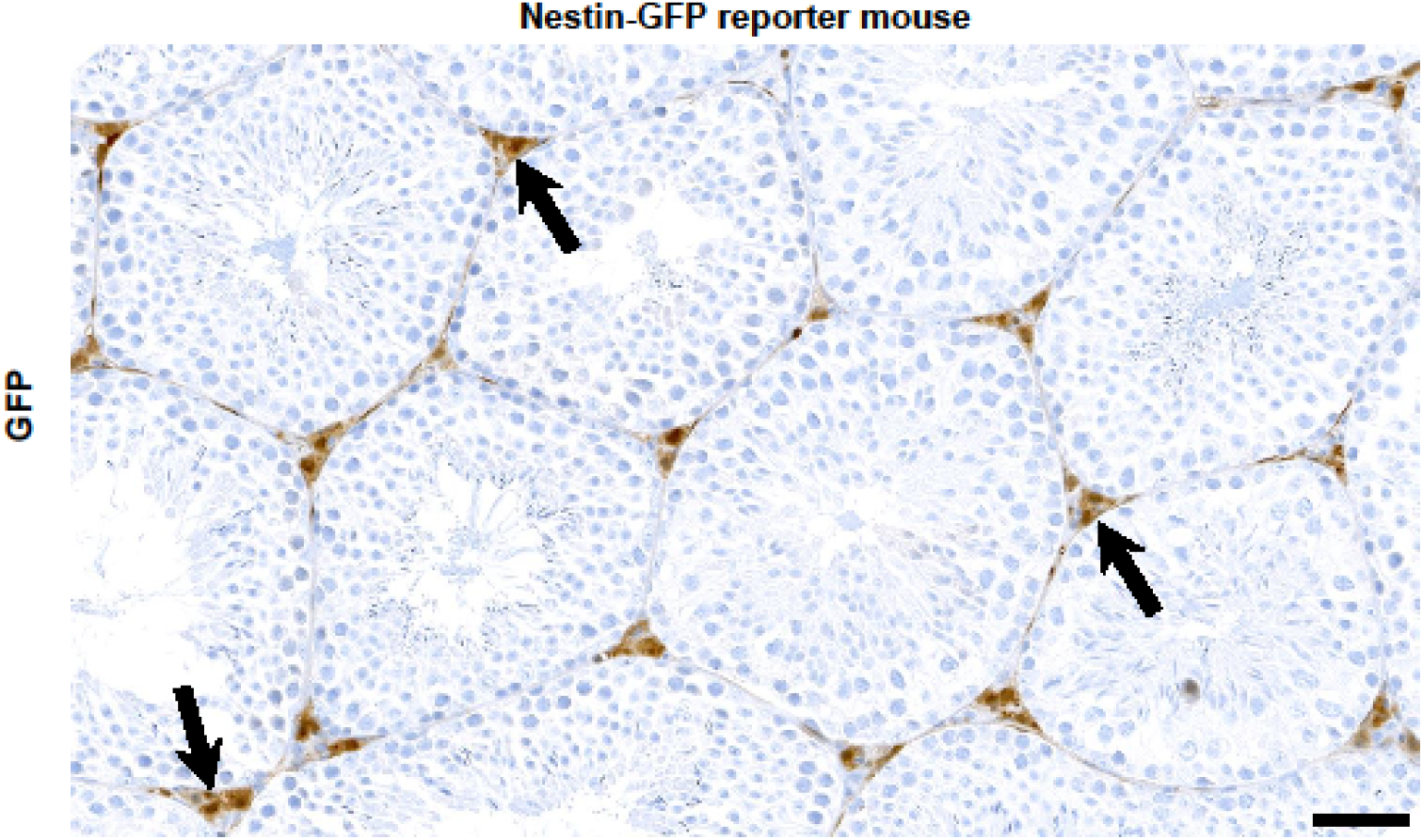
Nestin expression in Leydig cells. Immunohistochemistry for GFP identifies diffuse signal in the Leydig cell population (arrows) of reporter mice with transgenic GFP expression under the Nestin promoter (n=3). Scale bar, 50 μm.

**Figure 2-figure supplement 1.**
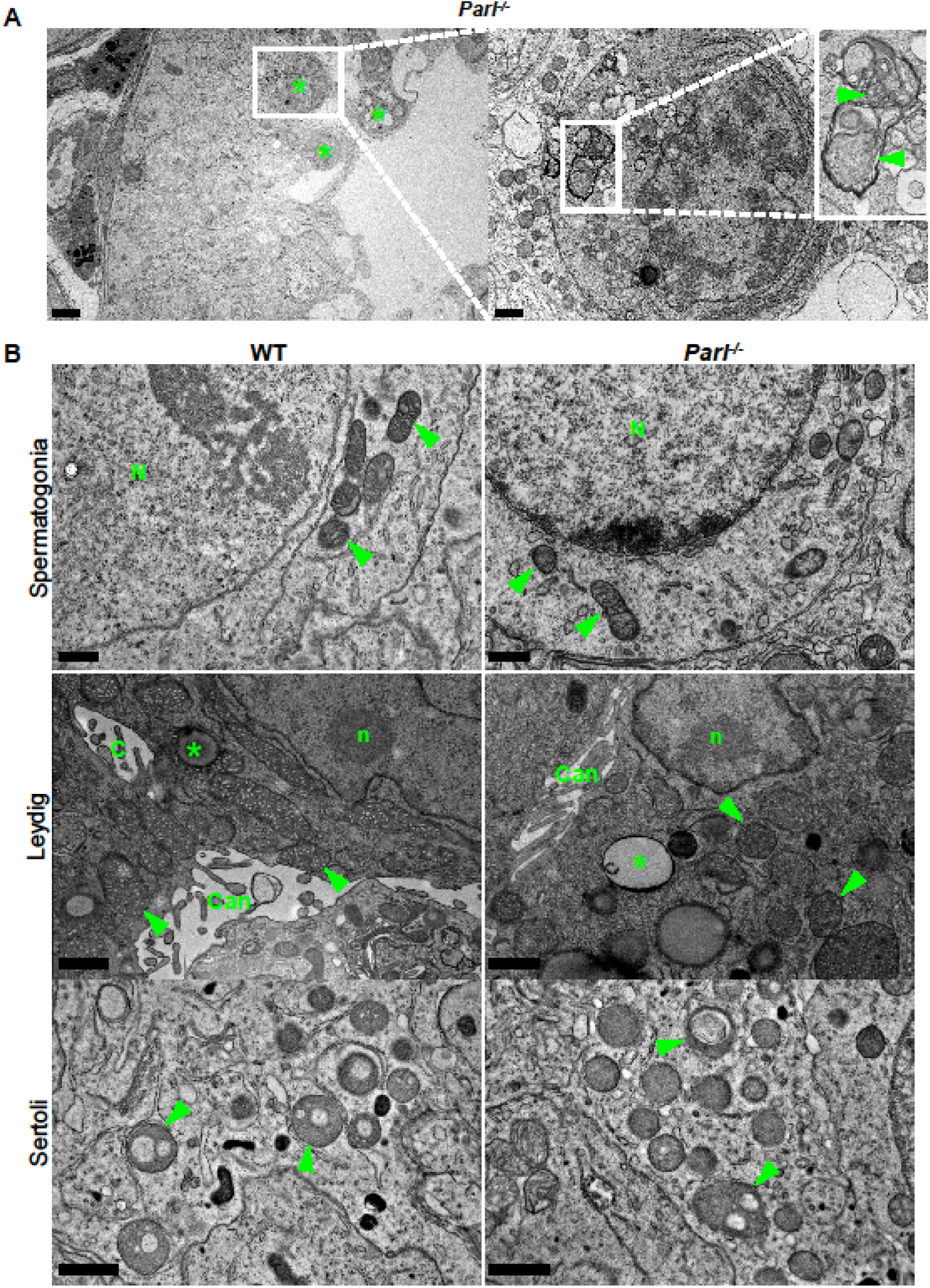
Mitochondrial and cellular ultrastructural abnormalities are restricted to arrested spermatocytes and absent in other testis cell types. (A) Degenerating/dying spermatocytes are mainly observed across the adluminal compartment of the seminiferous tubule (left panel, asterisks). At higher magnification (right panel), the cytoplasm of the degenerating spermatocyte shows multifocal cisternae distention and disruption of the endoplasmic reticulum with abundant accumulation of irregular coils of membranous material wrapped around damaged organelles including mitochondria (inset, arrowheads). Irregular nuclear infoldings and chromatin clumping are also evident (n=3). Scale bars, 5 μm (left panel) and 1 μm (right panel). (B) Ultrastructural abnormalities in PARL-deficient mice are not evident in spermatogonia, Leydig cells, and Sertoli cells. Spermatogonia (top panels) characterized by large round nuclei (N) and scant cytoplasm with scattered small oval mitochondria with lamellar cristae (arrowheads); scale bar, 0.5 μm. Leydig cells (middle panels) typically characterized by nuclei with a single prominent nucleolus (n), intercellular canaliculi with rudimentary microvillus processes (Can), large round to elongated mitochondria with dense tubular crista (arrowheads), and scattered cytoplasmic lipid droplets (asterisks); scale bar, 1 μm. Cytoplasmic projections of Sertoli cells (bottom panels) with typical round mitochondria characterized by few often dilated tubular cristae (arrowheads) (n=3). Scale bar, 0.5 μm.

**Figure 2-figure supplement 2.**
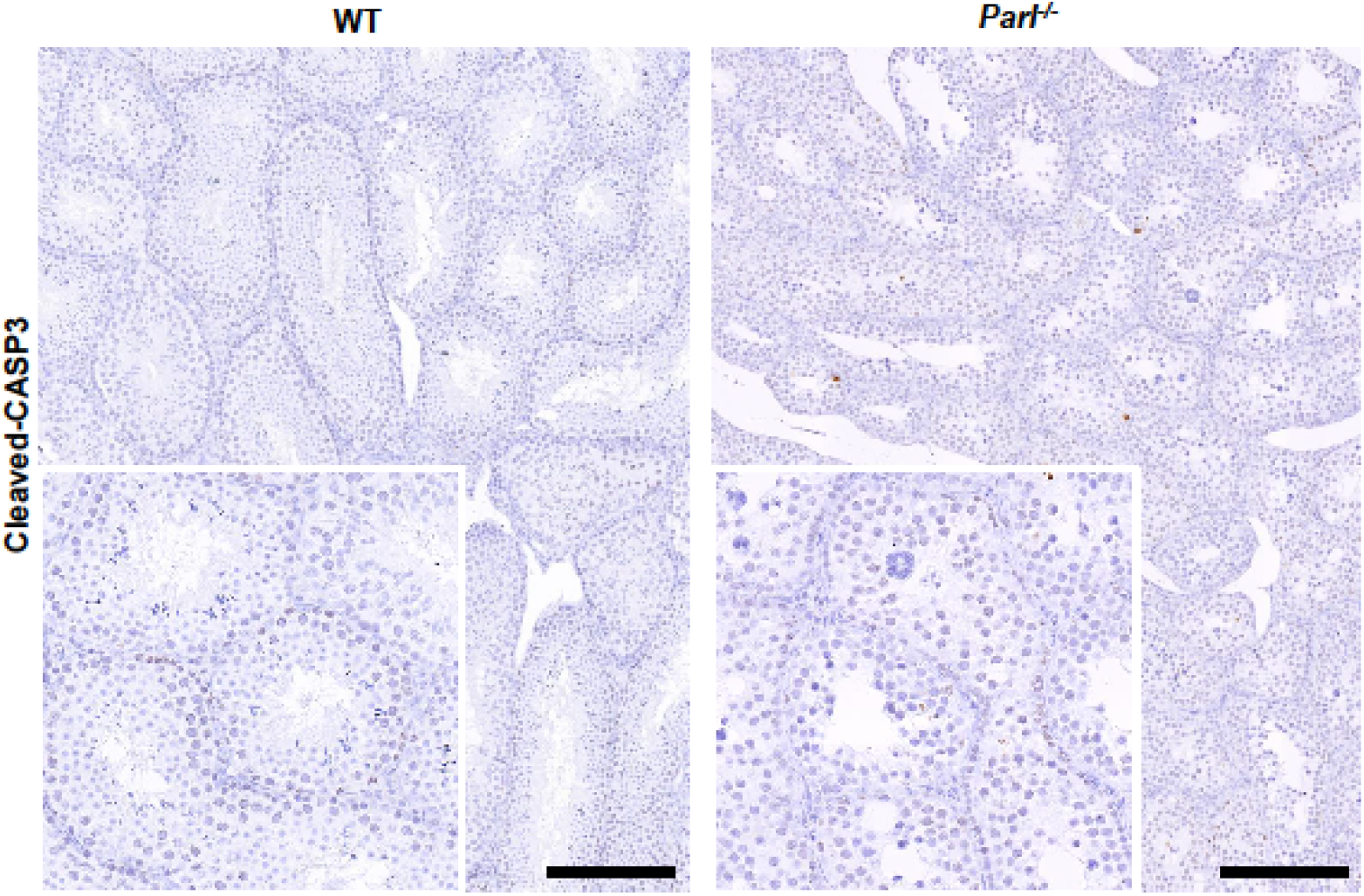
Absence of apoptosis in degenerated *Parl^-/-^* testis. Cleaved-Caspase-3 immunohistochemistry on testis from 7-week-old *Parl^-/-^* mice and WT littermates (n=3). The maturation defect and degenerative changes of PARL-deficient seminiferous tubules are not associated with the canonical activation of caspase-dependent apoptotic cell death as confirmed by the substantial lack of Caspase-3 cleavage. Scale bar, 200 μm.

**Figure 6-figure supplement 1.**
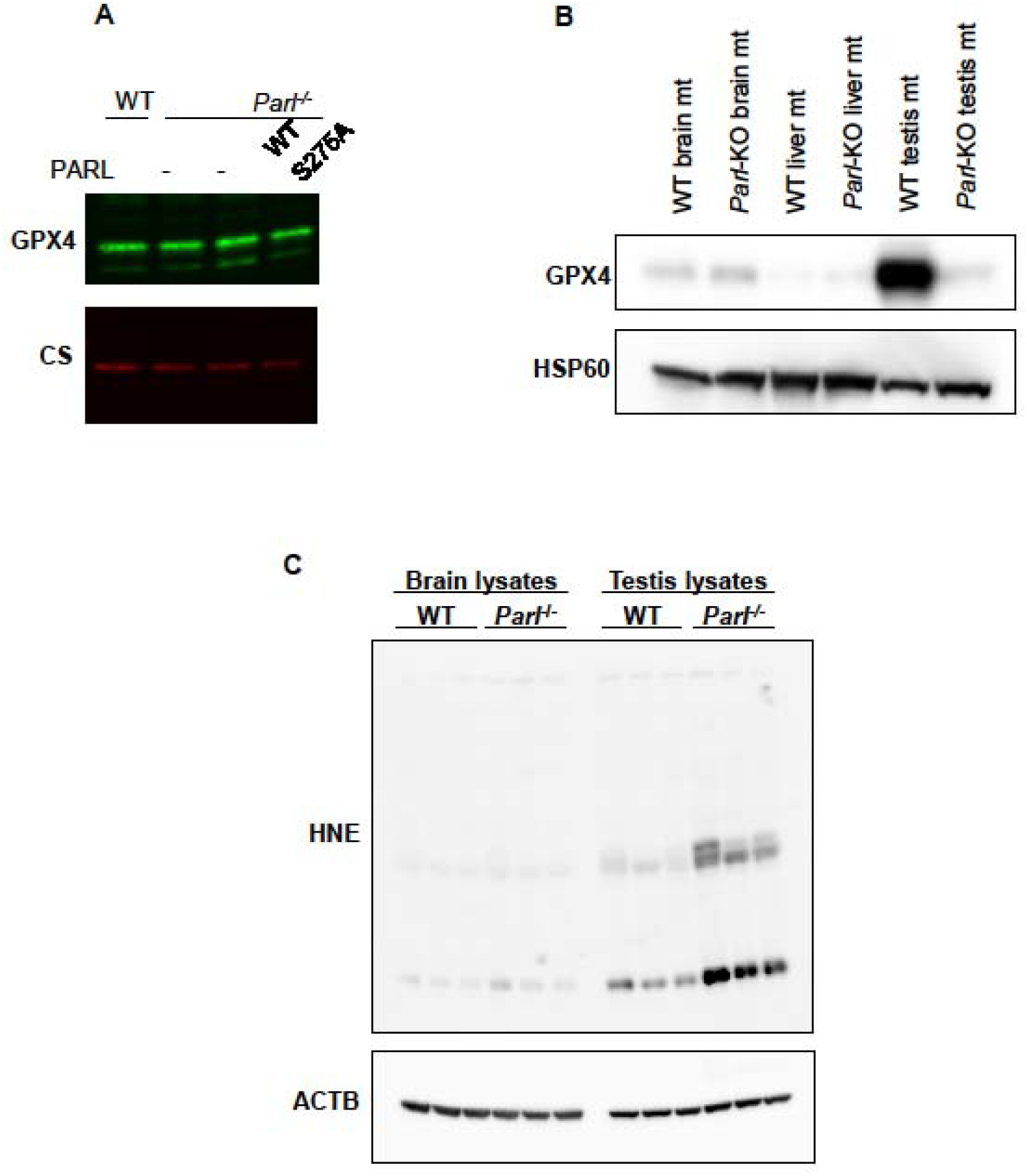
Lack of effect of PARL proteolytic activity on GPX4 expression in vitro and testis-specific induction of ferroptosis in PARL-deficient mice. (A) 20 μg of total protein from WT and *Parl^-/-^* MEFs complemented or not with WT or catalytic inactive PARL S275A were separated by SDS page and immunoblotted with GPX4 antibody. Citrate synthase (CS) is the loading control. (B) Mitochondria isolated from brain, liver and testis of 6-week-old WT and *Parl^-/-^* mice (n=3) were immunoblotted with antibodies for GPX4 and HSP60. HSP60 is the loading controls. GPX4 deficiency is evident in mitochondria isolated from *Parl^-/-^* testis, but not from other organs. (C) Brain and testis total lysates obtained from 6-week-old WT and *Parl^-/-^* mice (n=3) were immunoblotted with antibodies for HNE and ACTB. ACTB is as the loading control. In absence of PARL, lipid peroxidation is specifically increased in testis but not in the brain. **Figure 6-figure supplement 1 source data 1** Original images for panel A. **Figure 6-figure supplement 1 source data 2** Original images for panel B. **Figure 6-figure supplement 1 source data 3** Original images for panel C.

**Figure 6-figure supplement 2.**
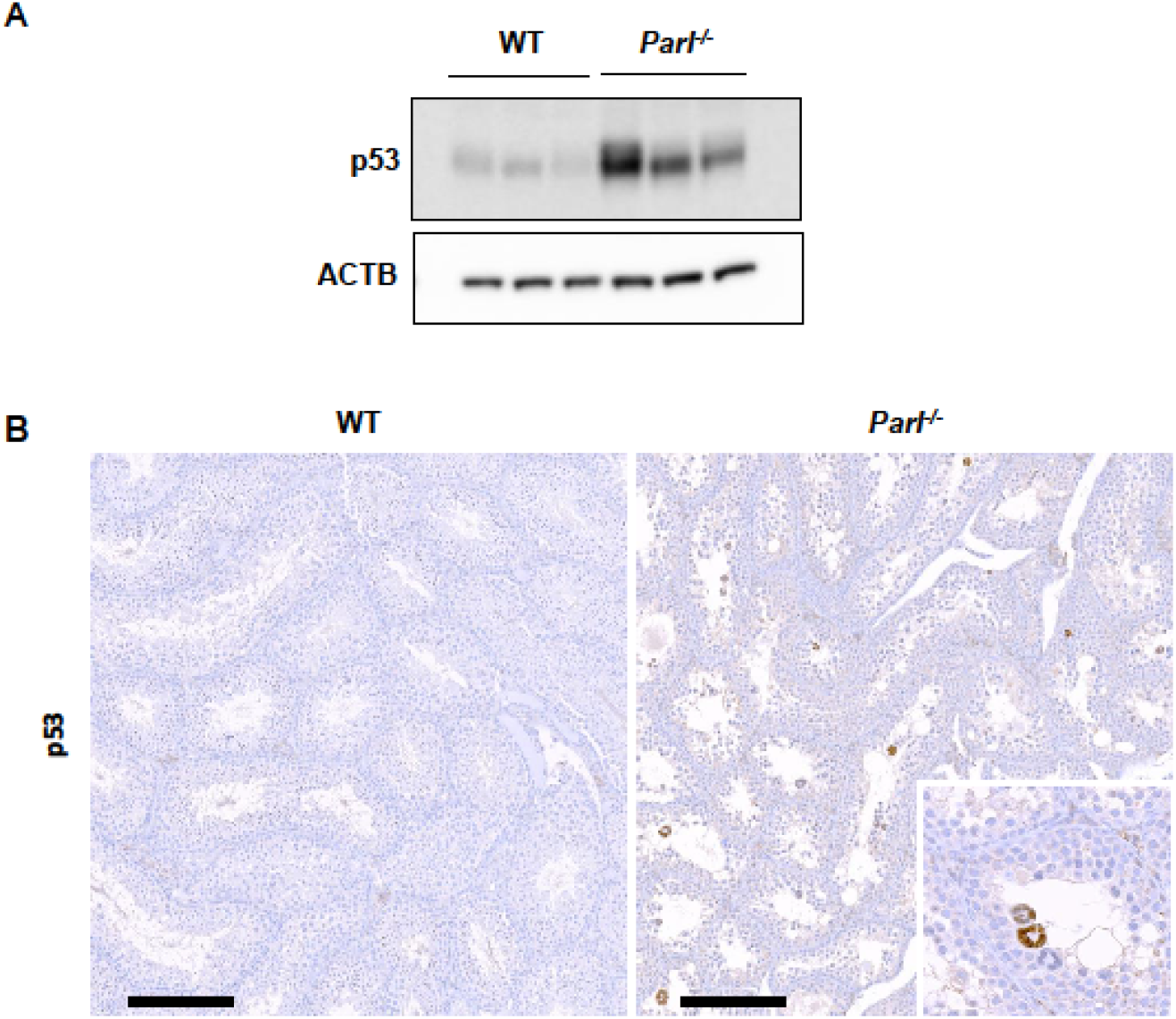
Increased activation of p53 in PARL-deficient spermatocytes undergoing ferroptosis. (A) Immunoblot analysis testis total lysates from 7 weeks-old WT and *Parl^-/-^* mice (n=3) with antibodies for p53 and ACTB. *Parl^-/-^* testes shows increased levels of p53 compared to WT littermates. (B) Immunohistochemical analysis confirms increased p53 expression in the seminiferous tubules of 7-week-old *Parl^-/-^* mice as compared to WT animals (n=3). Nuclear immunolabeling is mainly detectable in the adluminal and exfoliated multinucleated spermatocytes (inset) suggesting that p53 upregulation in *Parl^-/-^* testis takes place during the late stages of degeneration. No p53 expression is detectable via immunohistochemistry in WT littermates. Scale bars, 200 μm. **Figure 6-figure supplement 2 source data 1** Original images for panel A. **Figure 1-figure supplement 1** **Figure 2-figure supplement 1** **Figure 2-figure supplement 2** **Figure 6-figure supplement 1** **Figure 6-figure supplement 2** **Figure 3—source data 1** **Figure 4—source data 1** **Figure 5—source data 1** **Figure 6-figure supplement 1 source data 1** **Figure 6-figure supplement 1 source data 2** **Figure 6-figure supplement 1 source data 3** **Figure 6-figure supplement 2 source data 1**

